# An N-terminal amphipathic helix governs activity and conformational dynamics of Nramp metal transporters

**DOI:** 10.64898/2026.05.17.725737

**Authors:** Majid Jafari, Hongyan Zhao, Yao Zhang, Tianqi Wang, Kenneth M. Merz, Jian Hu

**Author notes:** Corresponding authors: Kenneth M. Merz, Jian Hu. Equally contributed to this work (Majid Jafari, Hongyan Zhao).

## Abstract

Nramps are a family of proton-coupled divalent metal ion transporters that play critical roles in maintaining homeostasis of essential metals. In humans, Nramp2 (DMT1) mediates iron uptake to support both cellular and systemic iron homeostasis, whereas Nramp1 exports metals from phagosomes, contributing to antimicrobial defense. A survey of experimentally solved Nramp structures reveals that an N-terminal helix (αA) immediately preceding TM1 is folded only in the outward-facing or outward-facing occluded states, raising the question of its role in Nramp transport. Here, we combined mutagenesis, transport assays, and molecular dynamics simulations to investigate αA. Our results show that deletion or substitution of conserved residues in αA markedly reduces Fe^2+^ transport. Simulations indicate that αA behaves as an amphipathic helix in the outward-facing state and preferentially stabilizes this conformation over the inward-facing state, thus acting as a key player in controlling conformational equilibrium. This study provides new insights into structural dynamics of Nramps and potentially sheds light on other LeuT-fold transporters.

## Introduction

Divalent metal transporter 1 (DMT1), also known as SLC11A2 or Nramp2, is a member of the natural resistance-associated macrophage protein (Nramp) family, a group of proton-coupled metal ion transporters conserved across kingdoms of life. In humans, DMT1 plays a central role in the uptake of ferrous iron (Fe^2+^), the bioavailable form of non-heme iron, across cell membranes. In the duodenum, DMT1 mediates dietary iron absorption into enterocytes following the reduction of ferric iron (Fe^3+^) to Fe^2+^.^1^ In the kidney, DMT1 is expressed in proximal tubules and contributes to the reabsorption of filtered iron, helping to prevent iron loss in urine.^2, 3, 4^ In peripheral tissues, DMT1 functions to facilitate endosomal iron release into the cytoplasm after endocytosis of transferrin-transferrin receptor complex, enabling iron utilization for essential processes such as heme synthesis and iron-sulfur cluster formation.^5^ Dysregulation of DMT1 has been linked to iron-related disorders. Loss-of-function mutations impair dietary non-heme iron absorption and disrupt endosomal iron trafficking, resulting in hypochromic microcytic anemia in rodents and humans.^5, 6, 7, 8, 9, 10, 11, 12^ Conversely, elevated DMT1 expression can contribute to pathological iron accumulation, as seen in hereditary hemochromatosis^13^ and neurodegenerative diseases such as Parkinson’s disease.^14^ Thus, a tight regulation of DMT1 is critical to maintain iron homeostasis.

Structural biology studies of Nramp family transporters have significantly advanced the understanding of the transport mechanism over the past decade. The first crystal structure was reported in 2014 for a bacterial Nramp homolog from *Staphylococcus capitis* (ScaNramp), revealing a LeuT fold with a core composed of ten transmembrane helices (TMs) that are arranged in a pair of inverted repeat and a metal binding site in the center of the transporter.^15^ Subsequent structures of other prokaryotic and plant Nramp homologs captured conformations in the inward-facing (IF), outward-facing (OF), and occluded (OC) states,^16, 17, 18, 19, 20, 21, 22^ establishing a rocking bundle model which is however different from the canonical mechanism for LeuT fold transporters. In this model, a unique set of moving TMs (TM1/5/6/10), rather than the helix bundle composed of TM1/2/6/7 for classical LeuT transporters,^23^ undergo significant displacement against the other relatively static TMs to alternately expose the transport site to either side of the membrane.^23, 24, 25, 26^ More recently, multiple cryo-electron microscopy (cryo-EM) structures of human DMT1 and its paralog Nramp1 in various states on the transport cycle were reported, extending the proposed rocking bundle model to human Nramps.^27^ Notably, in an outward-facing OC (OOC) state of Nramp1, a short N-terminal helix precedes TM1 (hereafter referred to as αA), as seen in the structures of DMT1 in the same conformational state, but this short segment is disordered in the IF state (**Figure 1A**). A survey of experimentally solved Nramp structures shows that αA can be folded in the OF state but disordered or unfolded in all structures in the IF states (**Figure 1B**). This strong preference suggests a dynamic role for αA during the transport cycle, leading to a hypothesis that it serves as a structural element that stabilizes or facilitates transitions between states. Bioinformatic analysis indicates that, although the αA sequence differs substantially across different kingdoms of life, it represents one of the most conserved regions for Nramps from metazoan (**Figure 1C**). However, the exact function of this short helix has not been clarified.

**Figure 1.**
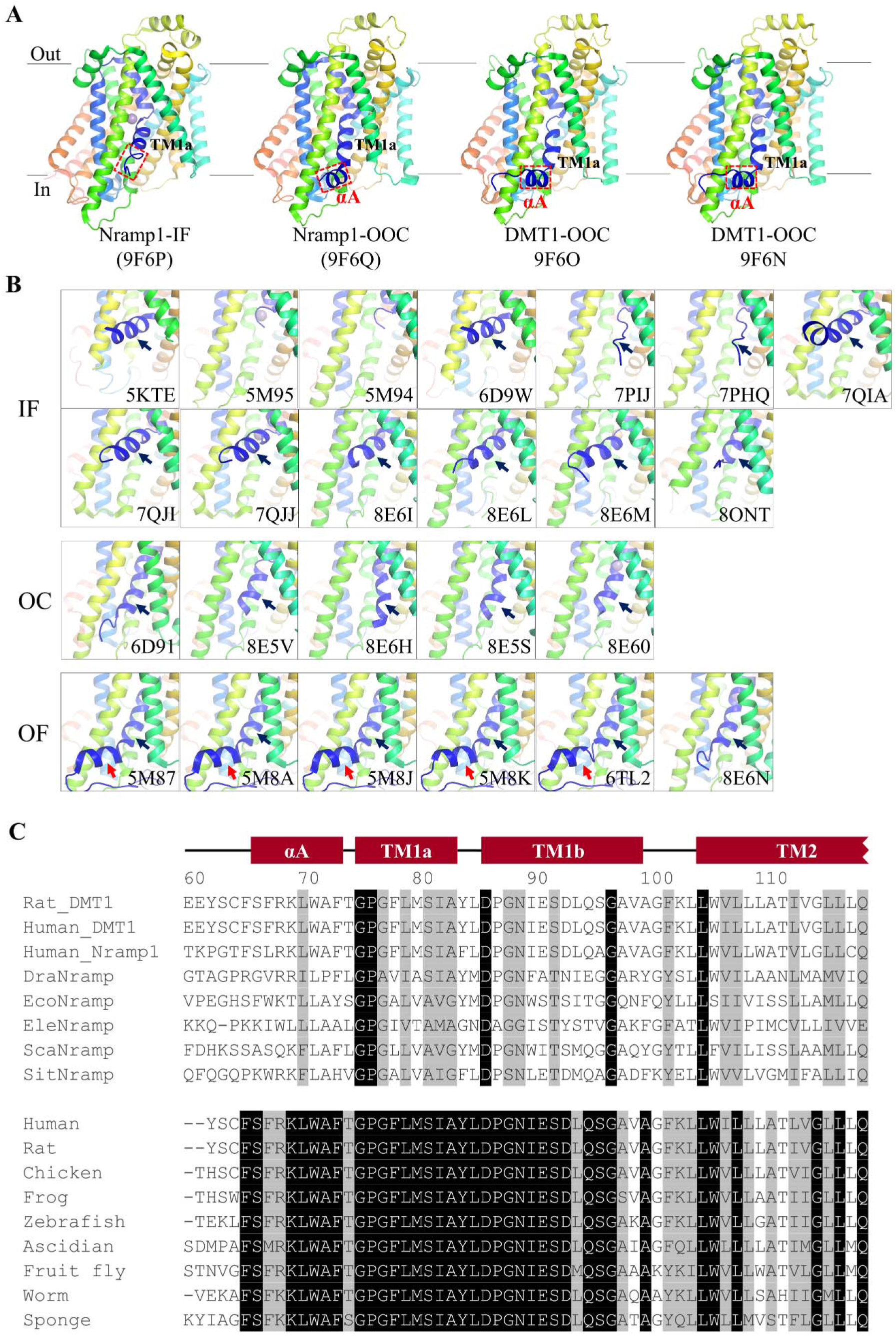
Structural comparison of Nramp structures. (**A**) Side-views structures of human DMT1 and Nramp1. αA and TM1a are labeled. The segment corresponding to αA is highlighted in the red dashed frame in each structure. OOC: outward-facing occluded state. (**B**) Comparison of the other Nramp structures in the region including TM1a (black arrow) and αA (red arrow). Structures are grouped based on the conformational states. (**C**) Comparison of Nramp sequences around TM1. The upper alignment includes the Nramps whose structures have been solved and the lower alignment compares the sequences of representative metazoan DMT1s. The red bars indicate α-helices according to the structure of human DMT1 (PDB: 9F6N). The residues are numbered based on the sequence of rat DMT1. The invariable residues and conserved residues are in black and grey backgrounds, respectively. The protein IDs in UniProt or NCBI are listed below. Rat_DMT1: *Rattus norvegicus*, O54902; Human_DMT1: Homo sapiens, P49281; Human_Nramp1: Homo sapiens, P49279; DraNramp: *Deinococcus radiodurans*, Q9RTP8; EcoNramp: *Eremococcus coleocola*, E4KPW4; EleNramp: *Eggerthella lenta*, C8WJ67; ScaNramp: *Staphylococcus capitis*, WP_002435436; SitNramp: *Setaria italic*, K3YRE7. Chicken_DMT1: *Gallus gallus*, B3F8C5; Frog_DMT1: *Xenopus laevis*, A0A974DS34; Zebra fish_DMT1: *Danio rerio*, Q1RLY4; Ascidian_DMT1: *Ciona intestinalis*, F6VMS4; Fruit fly_DMT1: *Drosophila melanogaster*, P49283; Worm_DMT1: *Caenorhabditis elegans*, Q21434; Sponge_DMT1: *Amphimedon queenslandica*, A0A1X7UN63.

In this work, we combined experimental and computational approaches to investigate the role of αA in transport activity and conformational dynamics. Deletion of αA or substitution of the residues in αA with alanine markedly reduced Fe^2+^ transport, and molecular dynamics (MD) simulations revealed that this amphipathic helix preferentially stabilizes the OF state over the IF state. Together, these findings support a model in which αA acts as a key determinant of Nramp function.

## Results

### Determination of DMT1 activity using a ^57^Fe transport assay

To study rat DMT1 (rDMT1), which shares 91.7% identical residues with the human counterpart and is a well-established prototypical family member with extensive biochemical characterization, we developed a cell-based stable iron isotope transport assay, adapted from an approach we used to study a zinc transporter.^28^ In brief, we transiently expressed rDMT1 in HEK293T cells and applied ^57^Fe, a stable iron isotope with a low natural abundance (2.1%), as the substrate in the transport assay for better quantification of iron transport activity (**Figure S1**). ^57^Fe in cells were then quantified by inductively coupled plasma mass spectrometry (ICP-MS), together with the phosphorous contents that was used to level out the cell number variation among samples. The resulting data showed that the cells expressing rDMT1 transported up to three folds more ^57^Fe than the cells transfected with an empty vector and ^57^Fe accumulation kept linearity up to 30 min incubation (**Figure S1A**). This iron transport activity is pH dependent with the optimal pH around 6.5 (**Figure S1B**). Applying a metal mixture to the cells expressing rDMT1 showed the transport activities, expressed as the molar ratios of metal and phosphorus, for Fe^2+^, Cd^2+^, Mn^2+^, Co^2+^, Ni^2+^, and Pb^2+^ but not for Zn^2+^ (**Figure S1C**). This result is consistent with early studies on mammalian DMT1s.^29, 30^

### αA is essential for DMT1 transport activity

To examine the role of αA in iron transport, we intended to delete the first 74 residues from the very N terminus of rDMT1 (**Figure 2A)**. Given that deletion of a chunk of N-terminal residues may affect protein folding and intracellular trafficking, it is necessary to determine the cell surface expression level of the variant and use it to calibrate the transport activity obtained from the cell-based transport assay. Guided by the AlphaFold predicted structure, we inserted a FLAG tag (DYKDDDDK) after G503 in the extracellular loop connecting TM11 and TM12, which is outside of the 10-TM core of the Nramp architecture. Functional study indicated that this modification led to a loss of iron transport activity by approximately 60% (**Figure 2B**, left) and the substantial activity allowed us to use this construct (hereafter referred to as exFLAG-WT) to examine protein expression on cell surface using immunofluorescence imaging and flow cytometry. As expected, the FLAG tag of exFLAG-WT was detected on the cell surface whereas another construct in which the FLAG tag was added at the C-terminus of rDMT1 (WT-C-FLAG) was not detectable when cells were not permeabilized in both flow cytometry and immunofluorescence imaging (**Figure 2B**, middle and right). Built on this construct, we generated two variants – Δ61 in which the first unstructured 61 residues were deleted and Δ74 in which αA and the preceding unstructured residues were all deleted. Using the ^57^Fe transport assay, we found that, when compared to exFLAG-WT, Δ61 showed a reduced activity by approximately 75% whereas Δ74 showed no activity (**Figure 2C**, left). Both variants were expressed, which was confirmed by confocal immunofluorescence imaging conducted on permeabilized cells (**Figure 2C**, middle), but their cell surface expression levels were reduced by approximately 50%, as quantified by flow cytometry (**Figure 2C**, right). Accordingly, the reduced activity of the Δ61 variant can be partially attributed to the reduced cell surface expression, but the completely abolished activity of the Δ74 variant and the only modestly reduced cell surface expression indicated that αA is essential for iron transport.

**Figure 2.**
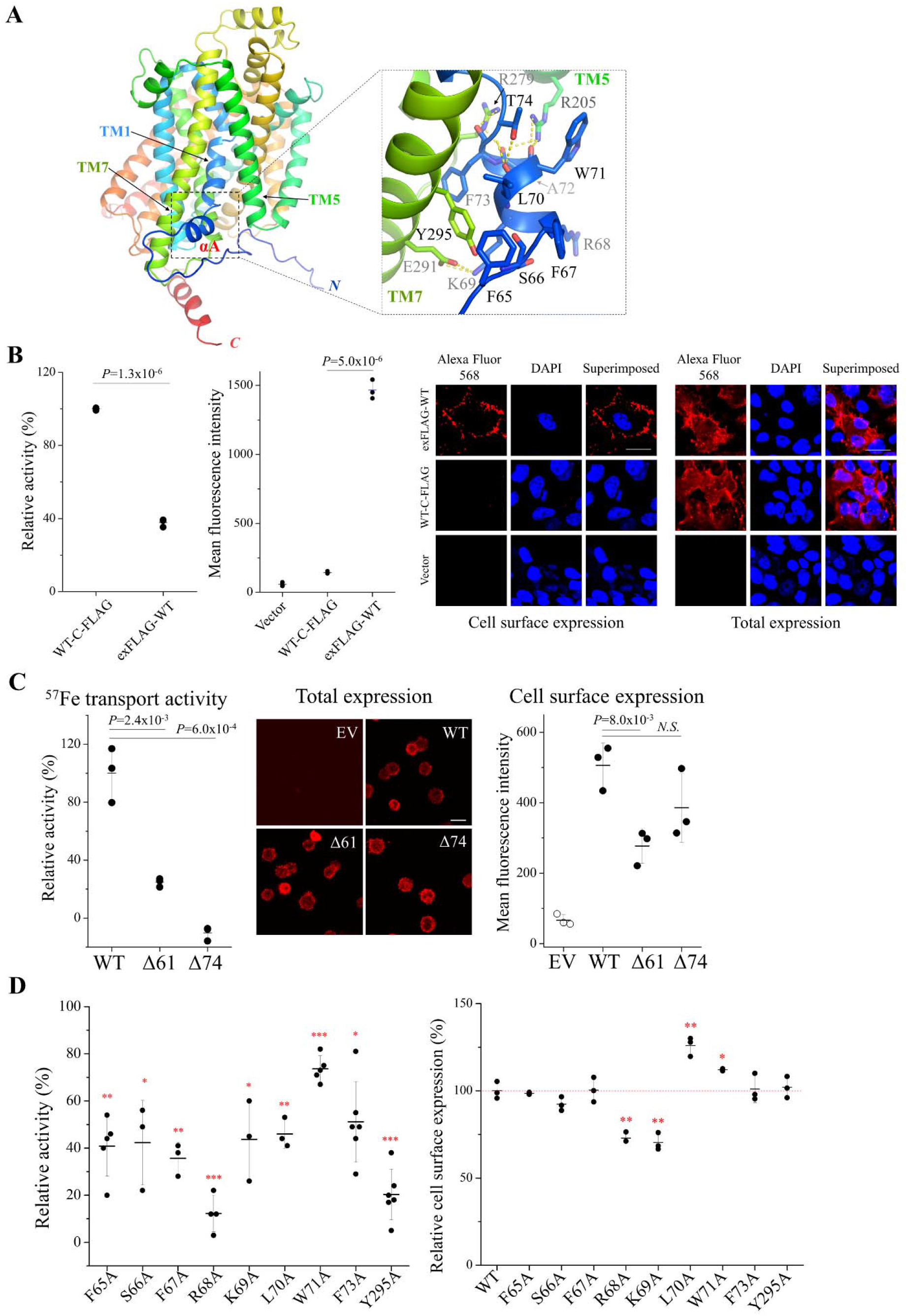
Functional characterization of αA in rDMT1. (**A**) Structural model of rDMT1 predicted by AlphaFold. The inset shows a zoomed-in view of αA and its polar interactions with the surrounding residues. The residues in αA and those involved in the interactions with αA are labeled and shown in stick mode. (**B**) Comparison of C-terminal FLAG tagged rDMT1 (WT-C-FLAG) and the construct with a FLAG tag inserted between the extracellular loop connecting TM11 and TM12 (exFLAG-WT). *Left*: Fe transport activity. The results shown are representative in 2-3 independent experiments; *Middle*: cell surface expressed detected by flow cytometry using an anti-FLAG antibody; *Right*: Immunofluorescence imaging. To detect cell surface expression level, cells were incubated with an anti-FLAG antibody without permeabilization, followed by staining with an Alexa Fluor 568-labeled secondary antibody. To detect total expression level, cells were permeabilized and then incubated with an anti-FLAG antibody and an Alexa Fluor 568-labeled secondary antibody. Cell nucleus were stained by DAPI. (**C**) Comparison of Fe^2+^ transport activities (*left*), total expression levels detected by immunofluorescence imaging (*middle*), and cell surface expression levels detected by flow cytometry (*right*) of empty vector (EV), exFLAG-WT (WT), the Δ61 and Δ74 variants. The results shown are from one representative experiment out of three independent experiments that produced similar results. (**D**) Characterization of the alanine-scanning variants. *Left*: Fe transport activity. Activities are expressed as the percentages of that of exFLAG-WT. The results shown are the averages from three to six independent experiments. One-sample two-tailed Student’s *t* test was performed to examine statistical significance. The *P* values from left to right are 0.002, 0.031, 0.0037, 0.0002, 0.029, 0.0044, 0.0242, 0.0009, and 0.0001, respectively. *Right*: cell surface expression level determined by flow cytometry. Cell surface expression levels are expressed as the percentages of that of exFLAG-WT. The results shown are from one representative experiment out of two independent experiments that produced similar results. Two-tailed Student’s *t* test was performed to examine statistical significance. The *P* values for R68A, K69A, L70A, and W71A are 0.0012, 0.0018, 0.0036, and 0.013, respectively. *: *P*<0.05; **: *P*<0.01; ***: *P*<0.001.

To examine the importance of individual residues in αA, we conducted alanine scanning on residues 65-73, covering from the highly conserved F65 to the last residue of αA (F73). Consistently with the high level of conservation of these residues among metazoans (**Figure 1C**), all alanine variants exhibited significantly reduced activity with R68A showing the greatest activity loss by nearly 90% (**Figure 2D**, left). Using flow cytometry, we quantified and compared the cell surface expression of all alanine variants and found that only two variants (R68A and L69A) showed modest reduction (**Figure 2D**, right). Accordingly, the diminished transport activities of these alanine variants are likely attributable to decreased transport efficiency rather than impaired expression or trafficking.

According to the structure of human DMT1 (PDB: 9F6O), these residues are involved in three types of interactions – F65, K69, L70, and F73 interact with TM7; F67, R68, and W71 potentially interact with the membrane lipids; and S66, as the helix-capping residue, stabilizes the helical structure of αA. The importance of the interaction of αA with TM7 was further confirmed by the Y295A variant, which loses activity by 80%. In fact, although the amino acid sequences of αA between bacterial Nramps and animal Nramps are barely conserved, the ways of αA to interact with TM7 are highly similar (**Figure 1B**), supporting a conserved role during evolution. As R68A loses the most activity among the alanine variants, the association of αA with the membrane and thus the position of αA relative to the membrane surface are likely crucial for the function of αA.

Overall, our biochemical data indicates that αA is essential for DMT1 transport activity. To elucidate the structural basis underlying the essential role of αA in Fe^2+^ transport by rDMT1, we performed MD simulations in the next section on human Nramp1 (hNramp1), whose structures in the IF and OOC states have been experimentally determined (**Figure 1A**).

### αA is a stable amphipathic helix in the OF state of hNramp1

Nramp1 and Nramp2 (or DMT1) share considerable sequence identity. For instance, the 10-TM core domains of human Nramps share 74% identical residues and their αA sequences are almost identical (**Figure S2**), suggesting that they employ the same or very similar transport mechanism. Human Nramps are the only animal Nramps whose structures have been solved. Nramp1 structure has been determined in two conformational states (IF and OOC) (**Figure 1A**), whereas Nramp2 structure was only captured in the OOC state. Therefore, we chose human Nramp1 in our simulation study.

We tested the structure stability of αA in hNramp1 in the OF and IF states using standard MD simulation. To generate a structural model of hNramp1 in the OF state, which has not been experimentally solved, we used AlphaFold to predict the OF state structure using the AF-Cluster approach.^31^ The generated model showed over 86% structural similarity to the previously solved *Eremococcus coleocola* Nramp in the OF state (PDB: 5M87, **Figure S3A**).^20^ The OF state of this Nramp homolog was used as reference because it has 12 TMs, whereas other Nramps with solved OF state structures do not. The OF state model of hNramp1 was then superimposed with the experimentally solved hNramp1 in the IF state (PDB: 9F6P) to reveal the TMs critically involved in the OF-IF interconversion (**Figure S3B**), and the pronounced movements of TM1/4/5/6/10 are consistent with the proposed transport mechanism for Nramps.^18, 26, 27^ The OF state model was also compared with the structure of hNramp1 in the OOC state (PDB: 9F6Q) (**Figure S3C**), revealing a highly superimposed αA/TM7 interface.

Then, we used the OF state model and the IF state structure to study the dynamics of αA in a lipid bilayer environment. In the OF state, αA behaves as an amphipathic helix and forms stably interactions with TM7, involving F53, K57, L58, F61, and Y283 (equivalent to F65, K69, L70, F73, and Y295 in rDMT1), and with membrane lipids where R56 and W59 (R68 and W71 in rDMT1, respectively) are at the lipid-water interface while F53, L55, and L58 (F65, F67, and L70 in rDMt1, respectively) face toward the hydrophobic core of lipid bilayers (**Figures 3A & 3B**). When the aromatic residues at the interface between αA and TM7 were substituted with alanine (F53A/F61A/Y283A), while αA maintained the helical structure during the simulations, its interactions with TM7 were severely disrupted (**Figure 3C**). To study the IF state, the missing αA was added to the experimentally solved structure (PDB: 9F6P) and the simulations were conducted under the same conditions as for the OF state. αA was partially unfolded in the initial model to study the effect of αA folding/refolding on the transporter’s IF state, but it became fully folded during the simulations. The result showed that the upward swing of TM1a forces αA to be buried deeper in the lipid bilayer and more tilted at the lipid-water interface (**Figures 3D, 3E & S3**). The unfavorable position in the bilayer and the largely reduced interactions with TM7 (**Figure 3F**) make αA more dynamic in the IF state than in the OF state, as indicated by the root mean square fluctuation (RMSF) analysis (**Figure 3G**), which explains why αA could not be experimentally resolved in the IF state.

**Figure 3.**
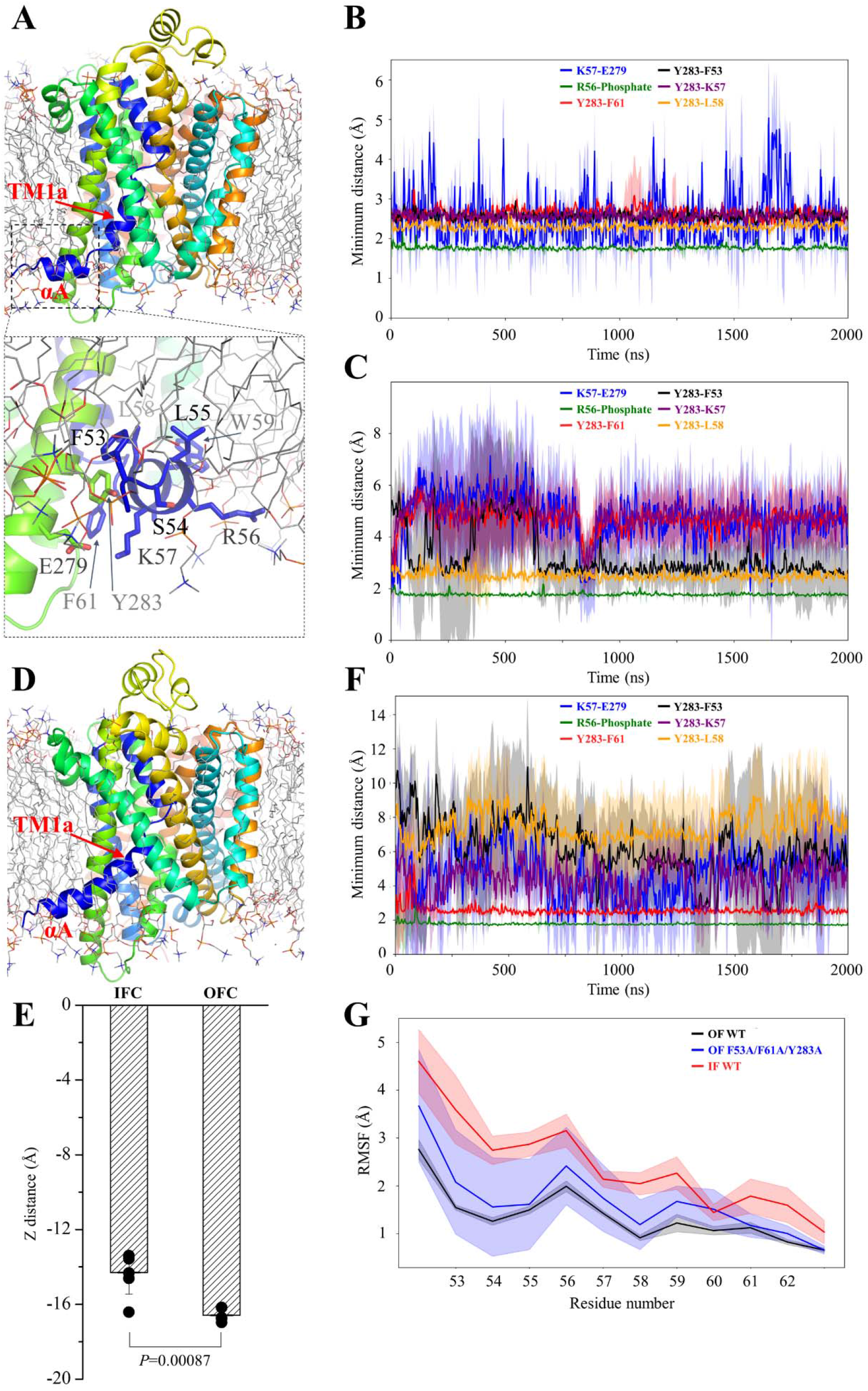
MD simulations of human Nramp1. (**A**) A representative snapshot in the MD simulations of the OF state. The protein is in rainbow color and the lipid molecules are depicted in the line mode with carbon, oxygen, phosphorus, and nitrogen in grey, red, orange, and blue respectively. The residues in αA and the contacting residues in TM7 are labeled in the stick mode in the zoomed-in view. (**B**) Distance changes of the indicated contacting pairs between αA and TM7 or between αA and membrane lipids in the OF state. (**C**) Distance changes of the indicated contacting pairs in the F53A/F61A/Y283A variant in the OF state. (**D**) A representative snapshot in the MD simulations of the IF state. (**E**) Embedment of αA in lipid bilayers in the OF and IF states. The average Z distance (vertical distance from the bilayer center) in each simulation is shown as one data point. A total of six independent simulations were conducted for each conformational state. Representative distance profiles in a single simulation are shown in **Figure S4**. (**F**) Distance changes of the indicated contacting pairs in the IF state with partially unfolded αA. (**G**) RMSF analysis of αA in the OF state (for wild type and the F53A/F61A/Y283A variant) and IF state (for wild type with added αA (labeled as IF WT)). For the simulation results in (B), (C), (F) and (G), the data represent the mean of three independent simulations, and the shaded regions indicate the standard deviation.

### αA preferentially stabilizes the OF state

Next, we investigated the role of αA in structural stability and dynamics of hNramp1 in the IF and OF states by comparing the wild-type protein with a Δ62 variant which corresponds to the Δ74 variant of rDMT1 in **Figure 2B**, and with the F53A/F61A/Y283A variant in which the interactions with TM7 were disrupted. In the OF state, deletion of αA led to a substantial increase in RMSF values for the residues throughout the sequence (**Figure 4A**), especially in the regions of TM4/5, TM9/10, and TM11/12. The triple variant also exhibited increased RMSF values in a pattern similar to that of the Δ62 variant, but to a lesser extent. In contrast, in the IF state, RMSF values were only modestly affected by αA’s positioning or the mutations at the interface with TM7 (**Figure 4B**).

**Figure 4.**
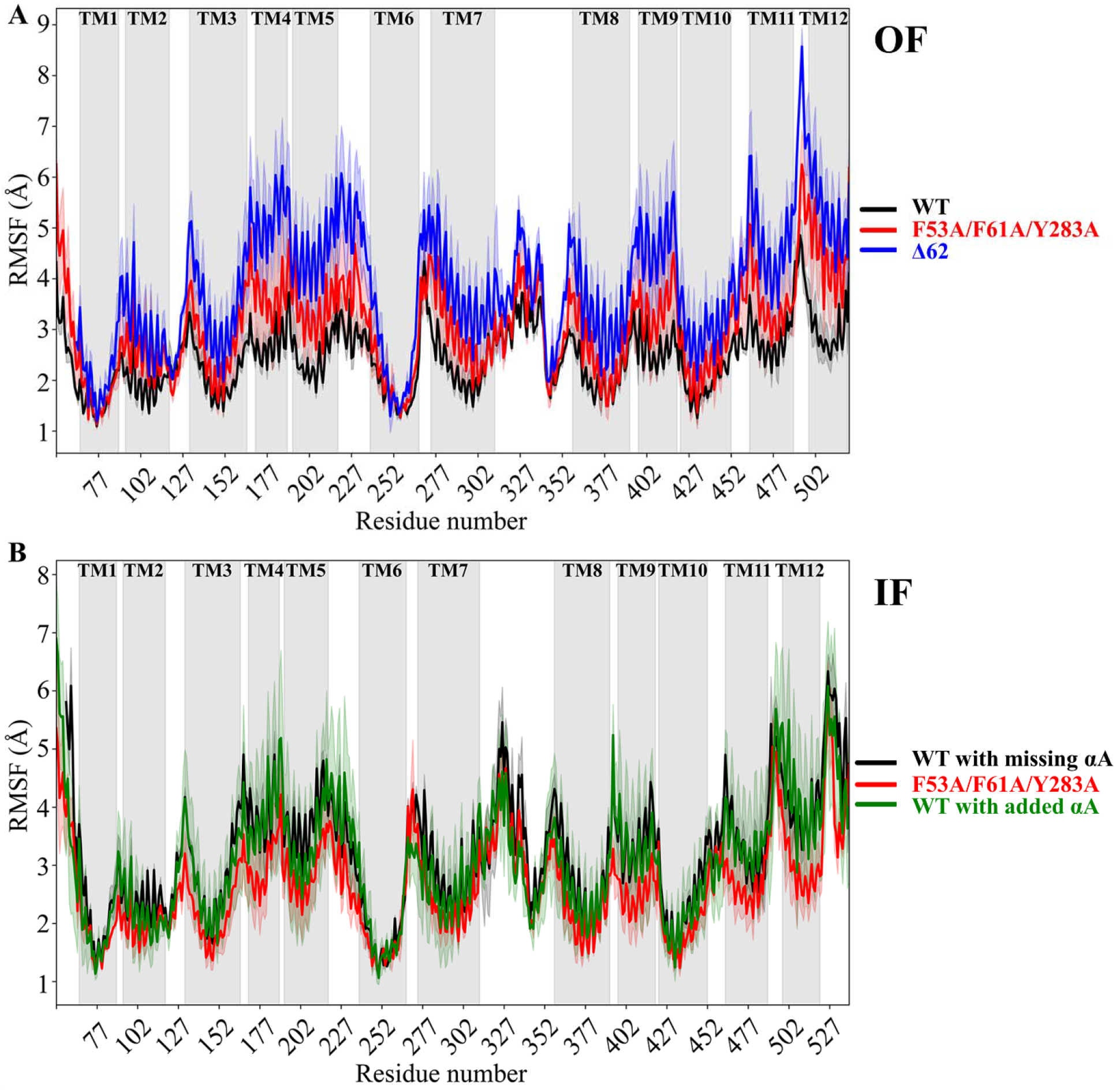
RMSF analysis of hNramp1. (**A**) hNramp1 in the OF state. WT (black) was generated using the AF-cluster method and hNramp1 structure (PDB: 9F6Q) as template. Mutations were introduced to this model to generate the variant model (red), and the first 62 N-terminal residues were deleted to generate the Δ62 variant model (blue). (**B**) hNramp1 in the IF state. The hNramp1 structure (PDB: 9F6P), in which αA is disordered and several residues are missing, was used directly in simulations (black). The predicted αA structure, with and without the indicated mutations, was added to this model (red and green, respectively). In both A and B, the X axis represents the residues numbers and the Y axis describes the movement of each residue measured in angstrom. The shaded regions with a light color represent the calculated standard deviations. The TMs are indicated by the grey areas. The results shown are averaged over three independent simulations, and the shaded regions indicate the standard deviation.

Significantly, although the OF state of the wild-type hNramp maintained a conformation that is 70-75% similar to the IF state, as calculated using the US-align method (**Figures 5A & 5E**),^32^ the Δ62 variant experienced a conformational rearrangement that increased the structural similarity to the IF state to 75-80% in the early stage of the simulations (**Figures 5A & 5D**). The triple variant showed the same trend, but again to a lesser extent (**Figures 5A & 5C**). When examining the structural snapshots, it was found that the averaged TM movement of the Δ62 variant in the OF state is reminiscent of the conformational switch from the OF state to the IF state (**Figures 5A & 5D, Video S1**), even though a complete switch to the IF state did not happen. In the control system, where the wild-type in the OF state was used as the starting model, we observed no significant movements in the TMs that are involved in the transporter’s major conformation change, except for a slight shift of TM10 (**Figure 5B**). Consistent with the result of the RMSF analysis, no significant change in similarity or structural rearrangement was observed in the IF state (**Figures 5E-H**), no matter whether a folded αA helix was included or missing. A modest shift in TM5 was observed in the systems containing αA (**Figures 5G & 5H**), suggesting a structural coupling between TM5 and αA. Overall, the MD simulation results indicate that αA, as an amphipathic helix that reversibly associated with membrane bilayer and TM7, preferentially stabilizes the OF state while exerting little influence on the IF state.

**Figure 5.**
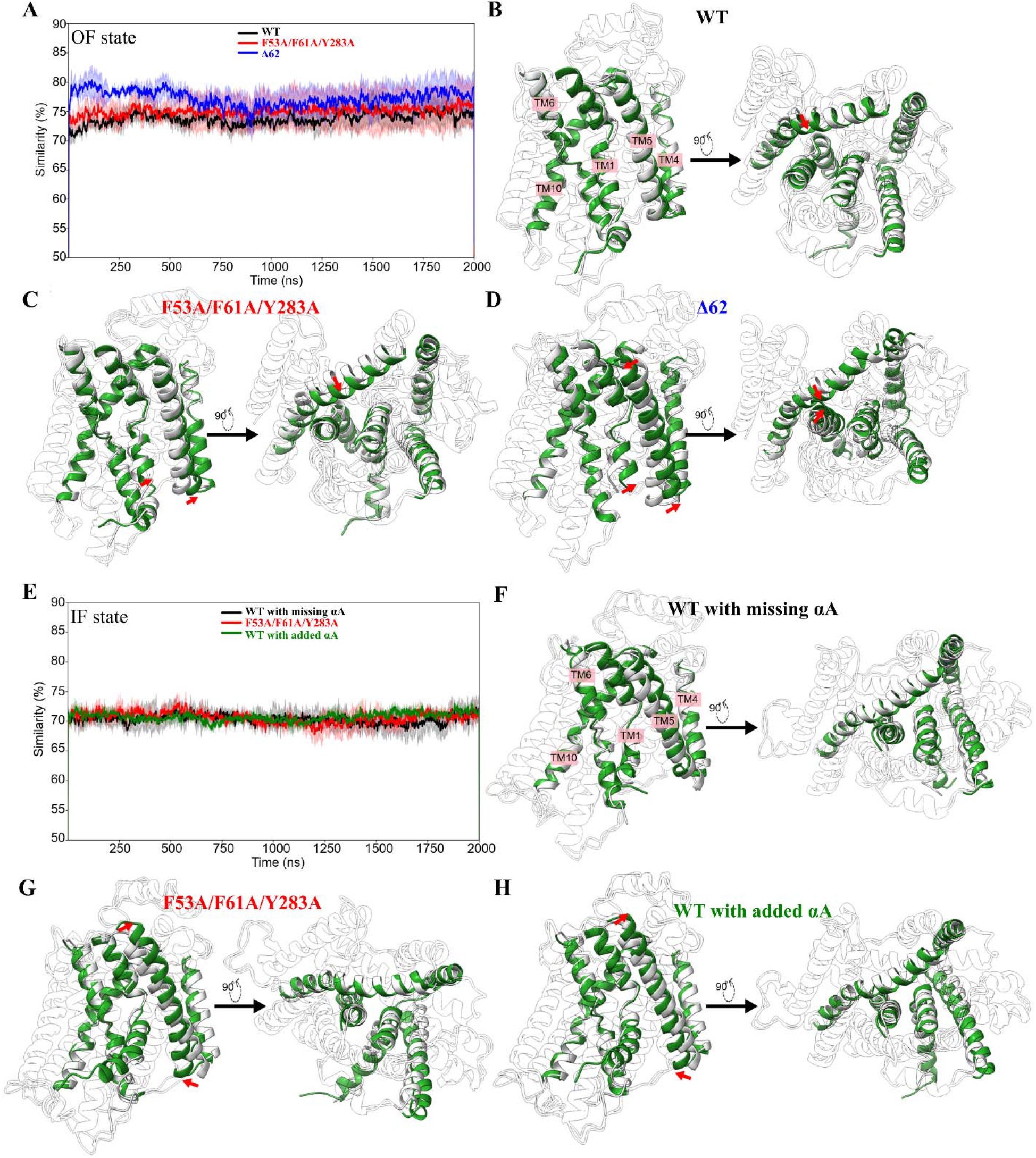
Structural similarity analysis of hNramp1 in MD simulations. (**A**) Changes in the structural similarity to the IF state during the simulations where the OF state models were used as the initial models. (**B-D**) Comparison of the cluster structure generated during the simulations using the initial models of the OF state of wild-type hNramp1, the F53A/F61A/Y283A variant, and the Δ62 variant (without αA), respectively. The cluster structures are shown in green and the initial model is in white. Only TM1/4/5/6/10 are shown for clarity. (**E**) Changes in the structural similarity to the OF state during the simulations where the IF state models were used as the initial models. (**F-H**) Comparison of the cluster structure generated during the simulations using the initial models of the IF state of wild-type hNramp1 (with missing αA), the hNramp1 with the partially unfolded αA carrying the F53A/F61A/Y283A mutations, and the hNramp1 with the partially unfolded αA carrying the wild-type residues, respectively. The cluster structures are shown in green and the initial model is in white. Only TM1/4/5/6/10 are shown for clarity. The pronounced structural changes are indicated by red arrows. The images show the averaged cluster structure from one representative simulation replicate.

## Discussion

Similar to other LeuT transporters, TM1 dynamics, especially the motion of cytoplasmic side of TM1 (i.e. TM1a), is a hallmark for the major conformational transition of Nramps. As shown in **Figure 1**, TM1a in a typical OF state is in a “down” position, sealing the cytoplasmic vestibule and blocking metal release from the transport site into the cytoplasm; in contrast, TM1a in the IF state swings to an “up” position, moving out of the helix bundle and thus creating an open vestibule that allows metal to be released. This significant conformational change is accompanied with and allowed by the coordinated movement of other TMs, in particular TM5, TM6, and TM10, as well as the pre-TM1 segment that forms a stable helix (i.e. αA) only in the OF or OOC state. This structural preference of the pre-TM1 segment raises the question whether αA plays a role in Nramp function or its helical structure is simply a consequence of the major conformational switch that creates a local environment to allow helix formation.

In this work, we applied combined experimental and computational approaches to address this question. Systematic mutagenesis and cell-based ^57^Fe transport assay revealed that αA is essential for transport function, as deletion of αA completely eliminated the iron transport activity while only modestly reduced the expression on cell surface (**Figure 2C**). The result of alanine scanning suggested that the interactions with both TM7 and membrane lipids contribute to the function of αA (**Figure 2D**). Using MD simulations, we demonstrated that αA behaves as an amphipathic helix stably associated with membrane surface in the OF state but exhibits greater structural fluctuation in the IF state (**Figure 3**). Importantly, we found that αA preferentially stabilizes the OF conformation (**Figure 4**), likely by anchoring TM1a via αA-TM7 and αA-lipid interactions, thereby biasing the transporter toward a substrate uptake-competent state (**Figure 5**). Based on these findings, we propose a model in which αA plays a key role in the Nramp transport cycle (**Figure 6**). In the metal-free OF state, the folded αA forms stable interactions with TM7 and the surrounding membrane lipids, thereby maintaining TM1a in a “down” position through a short linker in between. Upon binding of an extracellular (or intraluminal) metal ion to the high-affinity transport site located in the middle of TM1 and TM6, the transporter undergoes an OF-to-IF transition. Metal binding promotes an upward swing of TM1a, pulling αA deep into the membrane and disrupting its interactions with TM7, which results in αA unfolding. Subsequently, the bound metal ion is released into the cytoplasm through the now-open cytoplasmic vestibule. Loss of metal coordination in the transport site then triggers the IF-to-OF transition, during which TM1a swings downward to close the cytoplasmic vestibule. This movement drives the refolding of αA, and the folded αA in turn stabilizes TM1a in the “down” position through extensive interactions with TM7 and membrane lipids. Together, this proposal offers new mechanistic insight into the conformational dynamics of a Nramp protein. At the same time, we acknowledge that additional experimental evidence validating the altered conformational equilibrium resulting from αA disruption would be insightful. Beyond its contribution to conformational equilibrium, αA may also serve additional functional roles that are not captured by the current MD simulations. Because the folding and behavior of amphipathic helices are highly sensitive to the lipid environment,^33^ it is plausible that Nramp activity can be modulated by membrane lipid composition and physical properties. We further speculate that the marked sequence variation of αA among Nramps from prokaryotes, plants, and metazoans (**Figure 1C**) reflects evolutionary adaptation to their distinct membrane lipid compositions.^34, 35, 36^

**Figure 6.**
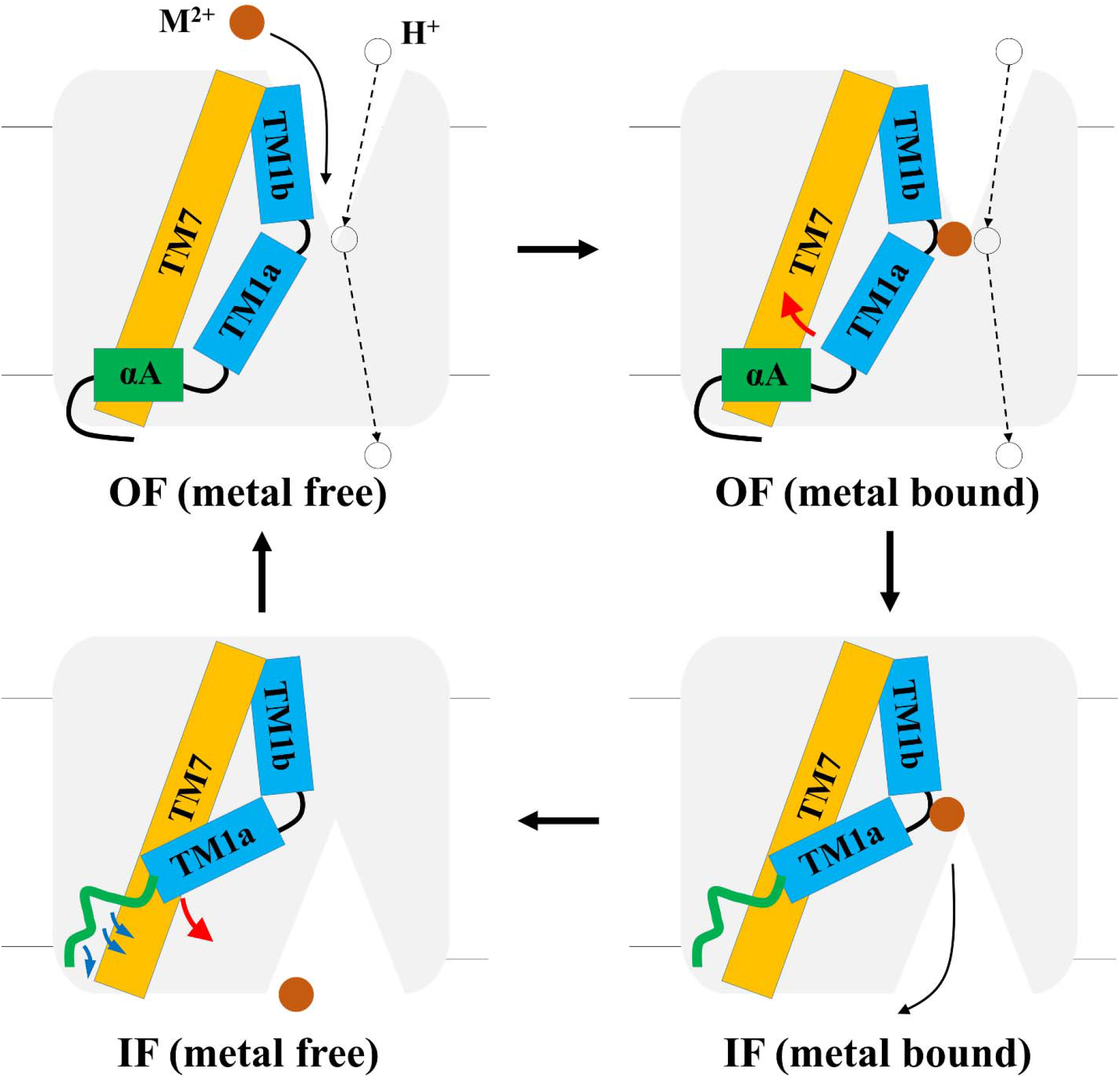
Proposed function of αA in controlling the dynamics of TM1a during the transport cycle of a Nramp transporter. In the metal-free OF state (upper right panel), the folded αA stabilizes the “down” position of TM1a. Metal binding to the transport site triggers the OF-to-IF conformational change in which TM1a undergoes an upward swing, likely accompanied with the unwinding of αA. After metal release to the other side of membrane (metal-bound IF state), TM1a swings back to the original position to block the vestibule (metal-free IF state), which is facilitated and stabilized by the interactions of αA with TM7 and membrane lipids (blue arrows). The transporter is depicted as a grey box and only TM1 and TM7 are shown as rectangles for clarity. A divalent metal ion is shown as a brown sphere and a proton is shown as an open sphere. The curved solid arrow indicates the metal translocation pathway. The dashed arrows indicate the separate proton translocation pathway.

The roles of pre-TM1 residues in conformational equilibrium have been reported for other LeuT-fold transporters. In the OF state of LeuT (PDB: 3TT1), the pre-TM1 residue W8 restrains TM1a dynamics through the interactions with the hydrophobic residues from the neighboring TMs,^37^ and the W8A mutation, which likely eliminates these interactions that stabilize the OF state, was found to favor an occluded IF state.^38^ In human serotonin transporter (SERT), mutations of the pre-TM1 residue T81 disrupt the conformational equilibrium and lead to a bias toward the IF state and thus abolishes the transport activity.^39^ However, neither W8 in LeuT nor T81 in SERT is in a helical structure or interacts with TM7. Thus, the use of an amphipathic helix to modulate conformational equilibrium represents a previously unrecognized mechanism for functional control in LeuT transporters. Indeed, amphipathic helices are well known to regulate membrane protein function,^40, 41, 42, 43^ and our finding provides a new paradigm for how such structural elements can control membrane protein activity.

## Materials and Methods

### Gene, plasmids, and reagents

The expression plasmid for rat DMT1 (UniProt ID: O54902) with a C-terminal FLAG tag was purchased from Sino Biological. The construct was modified by removing the C-terminal FLAG tag and inserting a FLAG-containing sequence (GS<u>DYKDDDDK</u>GS) after G503 in the linker between TM11 and TM12 (FLAG tag sequence is underlined). Mutations were generated using QuikChange® site-mutagenesis kit (Agilent Inc.) and verified by DNA sequencing. All the primer sequences for mutagenesis are listed in **Table S1**. Plasmids were purified using Miniprep (Promega, Cat#A1460) or Maxiprep (QIAGEN, Cat# 12163). ^57^FeCl_3_ was purchased from Sigma-Aldrich (Cat#790427, Lot#MBBD4771, 99.9% chemical purity and 95% isotope purity). ^57^FeCl_3_ was dissolved in 1 M HCl to the final concentration of 100 mM. Other reagents were purchased from Sigma-Aldrich or Thermo Fisher.

### Cell culture and transfection

HEK293T (ATCC, Cat#CRL-3216) were cultured in DMEM (Thermo Fisher, Cat#11965092) supplemented with 10% (v/v) FBS (Thermo Fisher, Cat#A5669801) and 1% antibiotic– antimycotic solution (Thermo Fisher, Cat#15240062) at 5% CO_2_ and 37 °C. Cells were seeded on polystyrene 24-well trays (Alkali Scientific, Cat#TPN1024) for 16 h and transfected with 800 ng DNA/well using polyethylenimine (Polysciences, Cat# 23966-2) in DMEM plus 10% FBS with a mass ratio of 1:3.

### Metal transport assay

48 hours post transfection, cells were washed with a washing buffer containing 20 mM 2-(N-morpholino)ethanesulfonic acid (MES), 142 mM NaCl, 5 mM KCl, 10 mM glucose, pH 6.0, followed by incubation with the buffer plus 1 mM freshly prepared ascorbic acid and 20 µM ^57^FeCl_3_ for 30 min at 37 °C. At the end of incubation, an equal volume of the ice-cold washing buffer containing 1 mM EDTA was added to the cells to terminate metal uptake, followed by three times of washing with ice-cold wash buffer before cell lysis in nitric acid.

### ICP-MS experiment

The ICP-MS experiments were conducted as described. All standards, blanks, and cell samples were prepared using trace metal grade nitric acid (70%, Fisher chemical, Cat#A509P212), ultrapure water (18.2 MΩ cm @ 25 °C), and metal free polypropylene conical tubes (Labcon, Inc.). For cell samples in polystyrene 24-well cell culture plates, 200 μl of 70% trace nitric acid was added to allow for initial sample digestion. Following digestion, 150 μl of the digested product was transferred into metal free 15 ml conical tubes. All cell samples were then incubated at 65 °C in a water bath for one hour followed by dilution to 5 ml using ultrapure water. The completed ICP-MS samples were analyzed using an Agilent 8900 Triple Quadrupole ICP-MS (Agilent Inc.) equipped with the Agilent SPS 4 autosampler, integrate sample introduction system, x-lens, and micromist nebulizer. Iron uptake by cells was expressed as the molar ratio of iron and phosphorus determined by ICP-MS, and iron transport activity was calculated by subtracting Fe uptake by the cells transfected with an empty vector from that by the cells expressing metal transporters.

### Flow cytometry

To quantify cell surface expression levels of FLAG-tagged rDMT1 (exFLAG-WT) and its variants using flow cytometry, resuspended cells after trypsin digestion were put on ice for 15 min, blocked by 2% bovine serum albumin (BSA), and then incubated with an anti-FLAG antibody (Sigma, Cat# F3165) at 4 μg/ml in Dulbecco’s phosphate buffered saline (DPBS, Gibco, Cat# 14190094) without Ca^2+^ and Mg^2+^ at 4 °C with gentle agitation for 1 h. Cells were then washed three times with DPBS buffer to remove unbound antibodies before incubation with Alexa Fluor 568-conjugated anti-mouse IgG (Invitrogen, Cat# A-11004) at a 1:400 dilution at 4 °C with gentle agitation for 2 h. After extensive wash with cold DPBS buffer, cells were fixed with 4% formaldehyde at room temperature for 15 min. Cell suspension in DPBS was subjected to flow cytometry analysis using a ThermoFisher Attune Cytpix flow cytometer. 10,000 cells were assessed for each condition. Cell debris was excluded based on FSC-A vs. SSC-A panel and single cells were selected through FSC-A vs. FSC-H gating. The mean fluorescence intensity (MFI) of each cell population was calculated. Data processing was performed in FlowJo 10.8.1 and GraphPad Prism 10.

### Immunofluorescence microscopy

To visualize cell surface expression, 24-well plates containing cell-seeded cover slides were placed on ice for 15 min to quench protein trafficking. After discarding culture medium, cells were washed once by using cold DPBS buffer and blocked by 2% BSA at 4 °C. Cells were then incubated with an anti-FLAG antibody at 4 μg/ml and 4 °C for 1 h. The cells were then washed three times by DPBS buffer and treated with Alexa Fluor 568-conjugated anti-mouse IgG diluted 1: 400 in DPBS buffer at 4 °C for 2 h. After washing three time, cells were fixed with 4% formaldehyde at room temperature for 15 min and washed three times before mounting on slides using fluoroshield mounting medium containing 4′,6-diamidino-2-phenylindole (DAPI, (Abcam, Cat# ab104139).

To visualize total expression, culture medium was discarded and cells (adherent or suspended) were washed once with DPBS buffer. Cells were then fixed using 4% formaldehyde at room temperature for 15 min. After washing three times with DPBS buffer, cells were permeabilized with 0.1% Tween-20 and blocked by 2% BSA in DPBS buffer at room temperature for 1 h. Cells were then incubated with an anti-FLAG antibody at 4 μg/ml and room temperature for 1 h. After washing three times with DPBS, cells were incubated with Alexa Fluor 568-conjugated anti-mouse IgG (Invitrogen, Cat# A-11004) diluted 1:400 in DPBS at room temperature for 2 h. Cells were then washed three times with DPBS, and coverslips were mounted onto slides using Fluoroshield mounting medium containing DAPI.

Images were taken with a 40 x objective using Nikon C2 confocal microscope.

### MD simulations

The coordinates of hNramp1 in the OCC and IF states were retrieved from the Protein Data Bank (PDB IDs 9F6Q and 9F6P, respectively). To generate the OF state structural model, the AF-Cluster method was applied under various parameter settings,^31^ including adjustments to the multiple sequence alignment (MSA) ratio, MSA depth, and the number of random seeds, both with and without structural templates. The crystal OF structure of *Eremococcus coleocola* Nramp (PDB: 5M87) and the hNramp1 Cryo-EM structure in its occluded state were used as templates. The best predictions were obtained when the 9F6Q structure was used as a template, with an MSA ratio of 8:16, random seed numbers of 10, 25, and 50, and MSA depths of 60000, 80000, and 100000. Across all these parameter combinations, as mentioned above, the resulting OF models showed a high degree of structural similarity (%86) to *Eremococcus coleocola* Nramp OF structure.

Six different simulation scenarios were designed to investigate the behavior of αA in human Nramp1, including three in the OF state and three in the IF state. The OF state structural model was generated as described above and was referred to as WT in **Figure 4A** (black profile). This structure was then used to generate the variant carrying the F53A/F61A/Y283A mutations, referred to as F53A/F61A/Y283A in **Figure 4A** (red profile). Next, the first 62 N-terminal residues were removed to create the Δ62 variant (blue profile in **Figure 4A**). Several key αA residues (F53, R56, and K57) that interact with TM7 and/or lipid molecules were missing from the IF state Cryo-EM structure (PDB: 9F6P), which was used as the starting coordinates for one scenario serving as a control (black profile in **Figure 4B**). The αA segment from the AlphaFold predicted model was added to the N-terminus of the hNramp1 Cryo-EM structure. The resulting model then underwent three rounds of energy minimization and equilibration to eliminate steric clashes. To model a partially unfolded αA, steered molecular dynamics (SMD) simulations were performed in which αA was gradually unfolded using a force constant of 1000 kcal·mol^-1^·Å^-2^ over a simulation time of 100 ns. The resulting structure was then used in MD simulations to generate the F53A/F61A/Y283A variant model (red profile in **Figure 4B**). The same structure, without mutations, was used to represent the wild type IF state (green profile in **Figure 4B**).

Protein structure models were embedded in a membrane model containing 200 1-palmitoyl-2-oleoyl-sn-glycero-3-phosphocholine (POPC) lipid molecules, with 100 lipids in each leaflet, using the CHARMM-GUI server.^44^ The systems were then solvated with TIP3P water molecules extending 25 Å on both the inner and outer sides of the membrane.^45^ Neutralization of the systems was performed by adding 0.15 M KCl to each system. To remove artificial clashes, three minimization phases were performed, each with positional restraints applied only to the protein and lipid molecules. The systems were then gradually equilibrated through six steps under constant number of particles, volume, and temperature (NVT) and constant number of particles, pressure, and temperature (NPT) conditions. During equilibration, the restraint strength was gradually reduced to allow safe and progressive equilibration. The duration of each equilibration step was 5 ns.

The equilibrated systems were then used for the production of MD phase. The temperature was maintained at 303 K using the Langevin dynamics thermostat, and the SHAKE algorithm was applied to constrain all bonds.^46^ Pressure was controlled at 1 bar using the Berendsen barostat with a relaxation time of 1 ps and semi-isotropic coupling.^47^ Long-range electrostatics were handled using the Particle Mesh Ewald method, and a cutoff of 9 Å was applied for nonbonded van der Waals and short-range electrostatic interactions. Local interactions between acidic residues and histidine residues were modeled using 12-6-4 LJ parameters.^48, 49, 50, 51^ Unbiased MD simulations were performed for 2 microseconds per system using the AMBER software package version 22,^52^ with the ff19SB^53^ and lipid21^54^ force fields. Three independent MD simulations, with extra runs carried out for certain cases as highlighted in the manuscript, were performed for each scenario.

Analysis of MD simulation results were conducted using custom Python scripts, CPPTRAJ,^55^ VMD,^56^ and UCSF Chimera.^57^ The structural alignment was performed on each frame of the MD simulations using US-align. The TM-score, which ranges from 0 to 1 (with 1 showing identical structures), was extracted for each frame and multiplied by 100 to represent structural similarity as a percentage throughout the simulations. For the simulations using the OF models as the initial models, the IF state was used as the reference, and *vice versa*.

### Statistics

We assumed a normal distribution of the samples and significant difference were examined using two-tail Student’s *t* test. Uncertainties shown in figures are reported as standard deviation (S.D.).

## Data availability

All raw and processed experimental and modeling data reported in the main text and SI are available upon request to Dr. Kenneth M. Merz and Dr. Jian Hu. All simulation data have been deposited in Zenodo (DOI: 10.5281/zenodo.17844557).

## Author contributions

J.H. and K.M.M. conceived the project and designed the study; H.Z., Y.Z., and T.W. conducted experiments; M.J. conducted MD simulations; All authors analyzed the data and wrote the manuscript.

## Acknowledgments

This work is supported by National Institutes of Health R35GM140931 (to J. H.) and R01GM130641 (to K.M.M.). The data presented herein were obtained using instrumentation in the MSU Flow Cytometry Core Facility. The facility is funded in part through the financial support of Michigan State University’s Office of Research & Innovation and Colleges of Osteopathic Medicine, Human Medicine, Veterinary Medicine, Natural Sciences, and Engineering. The Attune CytPix is supported by the Equipment Grants Program, award #2022-70410-38419, from the U.S. Department of Agriculture, National Institute of Food and Agriculture. The content is solely the responsibility of the authors and does not necessarily represent the official views of the National Institutes of Health.

