## Supplementary Information for "An N-terminal amphipathic helix governs activity and conformational dynamics of Nramp metal transporters"

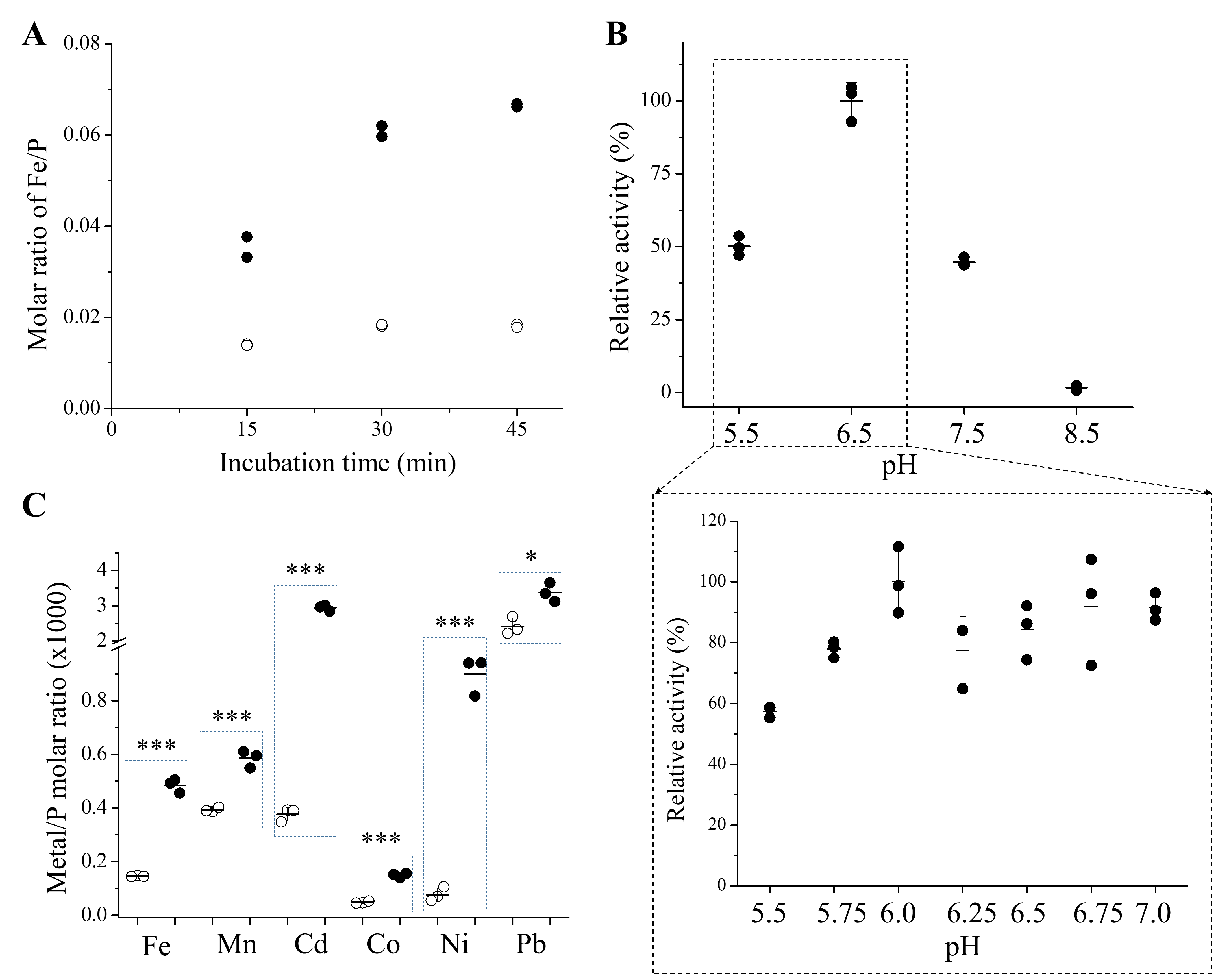

**Figure S1.** Functional characterization of rDMT1 using ^57^Fe and other non-radioactive metals. (**A**) Time courses of ^57^Fe uptake by HEK293T cells expressing rDMT1 (solid dots) and cells transfected with an empty vector (empty circle). (**B**) pH-dependent Fe transport activity. Solid dots represent the data of three replicates included for each pH. (**C**) Substrate specificity of rDMT1. In the cell-based transport assay, a metal mixture containing ^57^FeCl_3_ (5 µM), MnCl_2_ (15 µM), CdCl_2_ (10 µM), CoCl_2_ (5 µM), NiSO_4_ (5 µM), ZnCl_2_ (5 µM), Pb(NO_3_)_2_ (15 µM) in a buffer containing 20 mM MES, 142 mM NaCl, 5 mM KCl, and 10 mM glucose at pH 6.5, along with 1 mM freshly prepared ascorbic acid, was applied to cells. After incubation for 30 min at 37 °C, metal uptake was terminated by adding an ice-cold buffer containing 1 mM EDTA. After extensive wash, cells were lysed in nitric acid for ICP-MS analysis. The phosphorous content of the same sample was also determined by ICP-MS and used to level out cell number variation among samples. Accordingly, transport activity is expressed as the molar ratio of metal and phosphorus. For each metal included in the mixture, the metal/P ratio for cells expressing rDMT1 are shown as solid dots and those for cells transfected with an empty vector are shown as empty circles. No zinc transport activity was detected in this assay and therefore not shown in the plot. Student’s *t* test was used to examine statistical significance. *: *P*<0.01; ***: *P*<0.001.

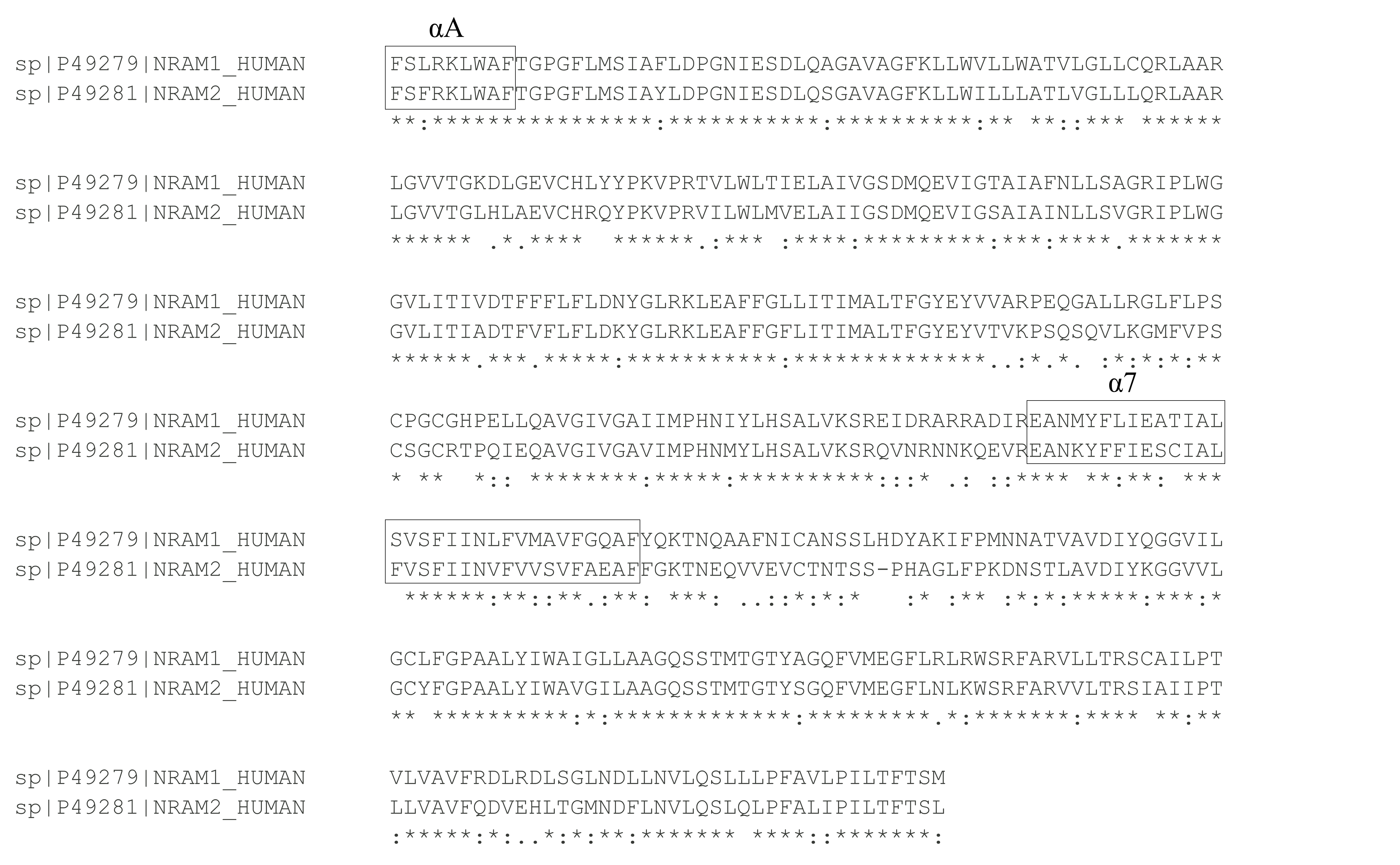

**Figure S2.** Sequence alignment of human Nramp1 and Nramp2. Only the 10-TM core domains are aligned. αA and TM7 are indicated by the frames.

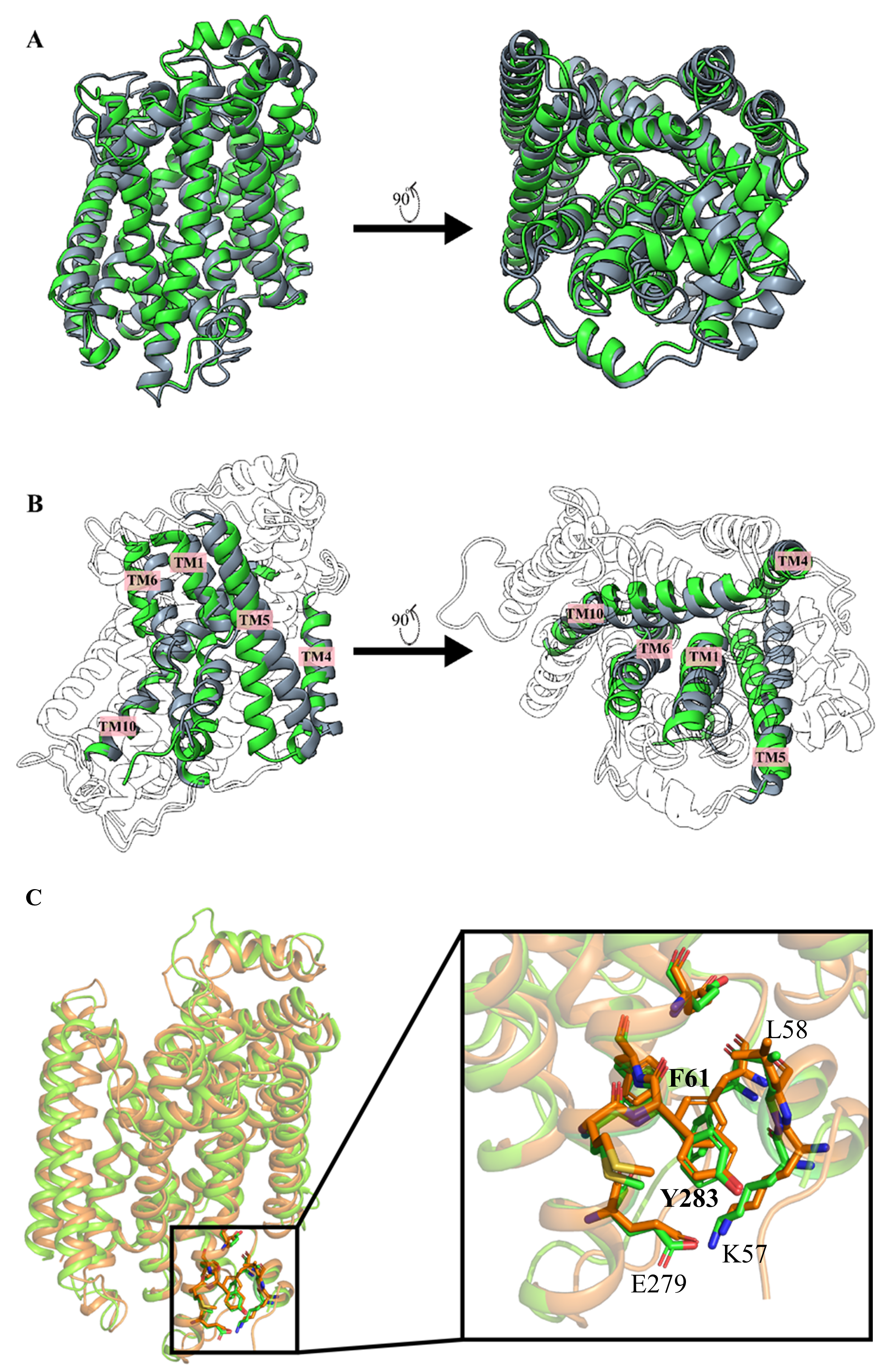

**Figure S3.** Structural comparison of AlphaFold-predicted hNramp1 in the OF state with experimentally solved structures. (**A**) Structure superimposition of the predicted hNramp1 in the OF state (green) and the *Eremococcus coleocola* Nramp in the OF state (gray, PDB: 5M87). (**B**) Structure superimposition of the predicted hNramp1 in the OF state (green) and the hNramp in the IF state (gray, PDB: 9F6P). Helices involved in the major conformational changes are colored, while the remaining helices are shown in transparent mode. (**C**) Superimposed structures of the predicted hNramp1 in the OF state (orange) onto the hNramp1 in the OOC state (green, PDB: 9F6Q). The αA-TM7 interface is highlighted in the zoomed-in view with interacting residues shown in stick mode. Note that F53 is not present in the OOC state structure.

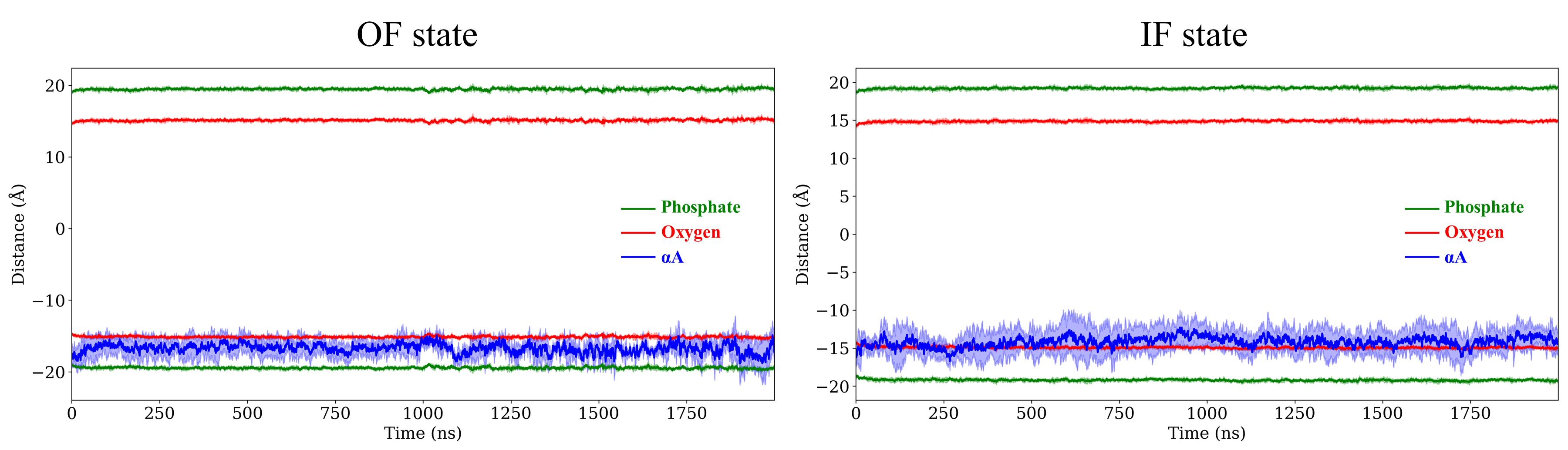
**Figure S4.** Representative profiles of the Z distance of αA in the OF (*left*) and IF (*right*) states during simulations. The Z distances of phosphate (green) and carbonyl oxygen of lipid molecules (red) are shown as references. Statistical analysis of six replicates for each condition is shown in **Figure 3E**.

**Table S1.** Primers used in this study.

| Primers | Sequences (5’-3’) |
| --- | --- |
| 67-73, All_Ala, forward | GAG GAG GAG TAC TCT TGT TTT AGT GCG GCG GCG GCG GCG GCG GCG ACA GGA CCT GGT TTT CTT ATG AGC |
| 67-73, All_Ala reverse | GCT CAT AAG AAA ACC AGG TCC TGT CGC CGC CGC CGC CGC CGC CGC ACT AAA ACA AGA GTA CTC CTC CTC |
| 67-73, All_Gly forward | GAG GAG GAG TAC TCT TGT TTT AGT GGC GGC GGC GGC GGC GGC GGC ACA GGA CCT GGT TTT CTT ATG AGC |
| 67-73, All_Gly reverse | GCT CAT AAG AAA ACC AGG TCC TGT GCC GCC GCC GCC GCC GCC GCC ACT AAA ACA AGA GTA CTC CTC CTC |
| Del 2-61 forward | GGG GCC GCC ACC AAG CTT GGT ACC ATG TAC TCT TGT TTT AGT TTC CGT AAA CTC |
| Del 2-61 reverse | GAG TTT ACG GAA ACT AAA ACA AGA GTA CAT GGT ACC AAG CTT GGT GGC GGC CCC |
| Del 2-74 forward | GGG GCC GCC ACC AAG CTT GGT ACC ATG GGA CCT GGT TTT CTT ATG AGC ATT GCC |
| Del 2-74 reverse | GGC AAT GCT CAT AAG AAA ACC AGG TCC CAT GGT ACC AAG CTT GGT GGC GGC CCC |
| GSDYKDDDDKGS insertion forward | GTC GTG  GTC TAC GTC CAG GAG CTA GGG  GGC AGC GAC TAC AAG GAC GAC GAC GAC AAG GGC AGC CAT GTG GCA CTG TAT GTG GTG GCT GCA |
| GSDYKDDDDKGS insertion reverse | TGC AGC CAC CAC ATA CAG TGC CAC ATG GCT GCC CTT GTC GTC GTC GTC CTT GTA GTC GCT GCC CCC TAG CTC CTG GAC GTA GAC CAC GAC |
| Del C-FLAG forward | GTA TCT AGA GGG GGT GGA GGC TAA GAT TAC AAG GAT GAC GAC GAT |
| Del C-FLAG reverse | ATC GTC GTC ATC CTT GTA ATC TTA GCC TCC ACC CCC TCT AGA TAC |
| F65A forward | CCT GAG GAG GAG TAC TCT TGT GCC AGT TTC CGT AAA CTC TGG GCC |
| F65A reverse | GGC CCA GAG TTT ACG GAA ACT GGC ACA AGA GTA CTC CTC CTC AGG |
| S66A forward | GAG GAG GAG TAC TCT TGT TTT GCC TTC CGT AAA CTC TGG GCC TTC |
| S66A reverse | GAA GGC CCA GAG TTT ACG GAA GGC AAA ACA AGA GTA CTC CTC CTC |
| F67A forward | GAG GAG TAC TCT TGT TTT AGT GCC CGT AAA CTC TGG GCC TTC ACA |
| F67A reverse | TGT GAA GGC CCA GAG TTT ACG GGC ACT AAA ACA AGA GTA CTC CTC |
| R68A forward | GAG TAC TCT TGT TTT AGT TTC GCC AAA CTC TGG GCC TTC ACA GGA |
| R68A reverse | TCC TGT GAA GGC CCA GAG TTT GGC GAA ACT AAA ACA AGA GTA CTC |
| K69A forward | TAC TCT TGT TTT AGT TTC CGT GCC CTC TGG GCC TTC ACA GGA CCT |
| K69A reverse | AGG TCC TGT GAA GGC CCA GAG GGC ACG GAA ACT AAA ACA AGA GTA |
| L70A forward | TCT TGT TTT AGT TTC CGT AAA GCC TGG GCC TTC ACA GGA CCT GGT |
| L70A reverse | ACC AGG TCC TGT GAA GGC CCA GGC TTT ACG GAA ACT AAA ACA AGA |
| W71A forward | TGT TTT AGT TTC CGT AAA CTC GCC GCC TTC ACA GGA CCT GGT TTT |
| W71A reverse | AAA ACC AGG TCC TGT GAA GGC GGC GAG TTT ACG GAA ACT AAA ACA |
| F73A forward | AGT TTC CGT AAA CTC TGG GCC GCC ACA GGA CCT GGT TTT CTT ATG |
| F73A reverse | CAT AAG AAA ACC AGG TCC TGT GGC GGC CCA GAG TTT ACG GAA ACT |
| Y295A forward | GAA GTT CGA GAA GCC AAT AAG GCG TTC TTC ATC GAG TCC TGC ATT |
| Y295A reverse | AAT GCA GGA CTC GAT GAA GAA CGC CTT ATT GGC TTC TCG AAC TTC |
